# Functionalized Mesoporous Silicas Direct Structural Polymorphism of Amyloid-β Fibrils

**DOI:** 10.1101/2020.01.13.904854

**Authors:** Michael J. Lucas, Henry S. Pan, Eric J. Verbeke, Lauren J. Webb, David W. Taylor, Benjamin K. Keitz

## Abstract

The aggregation of Amyloid-β (Aβ) is associated with the onset of Alzheimer’s Disease (AD) and involves a complex kinetic pathway as monomers self-assemble into fibrils. A central feature of amyloid fibrils is the existence of multiple structural polymorphs, which complicates the development of disease-relevant structure-function relationships. Developing these relationships requires new methods to control fibril structure. In this work, we demonstrate that mesoporous silicas (SBA-15) functionalized with hydrophobic (SBA-PFDTS) and hydrophilic groups (SBA-PEG) direct the aggregation kinetics and resulting structure of Aβ_1-40_ fibrils. The hydrophilic SBA-PEG had little effect on amyloid kinetics while as-synthesized and hydrophobic SBA-PFDTS accelerated aggregation kinetics. Subsequently, we quantified the relative population of fibril structures formed in the presence of each material using electron microscopy. Fibrils formed from Aβ_1-40_ exposed to SBA-PEG were structurally similar to control fibrils. In contrast, Aβ_1-40_ incubated with SBA-15 or SBA-PFDTS formed fibrils with shorter cross-over distances that were more structurally representative of fibrils found in AD patient-derived samples. Overall, these results suggest that mesoporous silicas and other exogenous materials are promising scaffolds for the *de novo* production of specific fibril polymorphs of Aβ_1-40_ and other amyloidogenic proteins.

**Significance Statement:** A major challenge in understanding the progression of Alzheimer’s Disease lies in the various fibril structures, or polymorphs, adopted by Amyloid-β (Aβ). Heterogenous fibril populations may be responsible for different disease phenotypes and growing evidence suggests that Aβ fibrils formed *in vitro* are structurally distinct from patient-derived fibrils. To help bridge this gap, we used surface-functionalized mesoporous silicas to influence the formation of Aβ_1-40_ fibrils and evaluated the distribution of resulting fibril polymorphs using electron microscopy (EM). We found that silicas modified with hydrophobic surfaces resulted in fibril populations with shorter cross-over distances that are more representative of Aβ fibrils observed *ex vivo*. Overall, our results indicate that mesoporous silicas may be leveraged for the production of specific Aβ polymorphs.

## Introduction

The aggregation of amyloidogenic proteins is associated with a number of neurodegenerative diseases, including Alzheimer’s Disease (AD)(1–3). Specifically, Amyloid-β (Aβ) aggregates and neurofibrillary tangles of tau are implicated in the progression of AD(4, 5). A key challenge in unraveling the pathology of AD and designing effective therapeutics is the paucity of structurefunction relationships for amyloid oligomers and fibrils. Unfortunately, the development of such relationships is complicated by the presence of multiple protein isoforms(6), post-translational modifications(7–9), and the range of potential fibril structures(10).

A central feature of amyloid fibrils is structural polymorphism(11). Fibril polymorphs differ in the registry, orientation (head-to head *versus* head-to-tail), and facial symmetry of the protein subunits. Consequently, a vast array of structures are theoretically possible(10). The structural differences between amyloid fibrils are important because each fibril polymorph potentially exhibits distinct disease- and other amyloid-controlled phenotypes(12–14). For example, the structural heterogeneity of Aβ fibrils may contribute to different AD subtypes and complicates therapeutic development(15). Thus, the structural characterization of different amyloid fibril polymorphs and an improved understanding of their biological role are critical steps in evaluating their influence on disease progression.

Amongst *in vitro* fibrils assembled from a single protein (e.g., Aβ_1-40_), a number of fibril polymorphs have been structurally characterized using X-ray crystallography, NMR spectroscopy, and electron microscopy (EM)(16–20). Additionally, recent electron microscopy studies have characterized specific fibril polymorphs of a-synuclein and Aβ from Parkinson’s and AD patients, respectively(21, 22). One notable finding from these studies was the presence of multiple structural polymorphs among *ex vivo* Aβ amyloid fibrils. More importantly, several studies have shown that Aβ fibrils prepared *in vitro* are structurally distinct from recent examples of Aβ fibrils isolated from AD patients(21, 22). Specifically, *ex vivo* fibrils displayed right-handed twists with hydrophobic residues exposed to solvent, while fibrils formed *in vitro* exhibited left-handed twists with a hydrophobic core(22). Given these differences, it is unclear what percentage of theoretical fibril structures are accessible *in vivo* or how aggregation conditions influence fibril structure(23–25). Overall, there is a pressing need for methods that enable the production or isolation of specific amyloid polymorphs as well as imaging techniques that can characterize heterogeneous populations of fibril structures.

Amyloid formation is a nucleation driven process heavily influenced by the surrounding environment(26–29). For example, a variety of environmental stimuli, including metal ions, chaperones, membranes, and other cellular machinery strongly impact amyloid formation(30–35). Although amyloid nucleation and growth is very complex, classical nucleation theory predicts that the environment of the critical nucleus should influence the structure of the resulting aggregate (Figure 1a)(36–38). Nucleation theory is widely applied to direct the crystallization of small molecule polymorphs, and nucleation driven control over amyloid structure has been used to produce fibrils of sufficient purity for X-ray crystallography and solid-state NMR spectroscopy through repeated seeding(17, 39). In addition to amyloid seeds, prior work has shown that exogenous materials, including nanoparticles, polymers, and small molecules can act as nucleants that influence the kinetics of amyloid aggregation(40–44). While the effect of these agents on the kinetics of amyloid formation has been widely studied, it is generally unclear how changing aggregation conditions dictates the structure of the resulting amyloid fibrils. Given these prior results and the potential influence over the nucleation step, we hypothesized that exogenous materials could affect the structure of Aβ fibrils and potentially direct the formation of specific fibril polymorphs.

**Figure 1.**
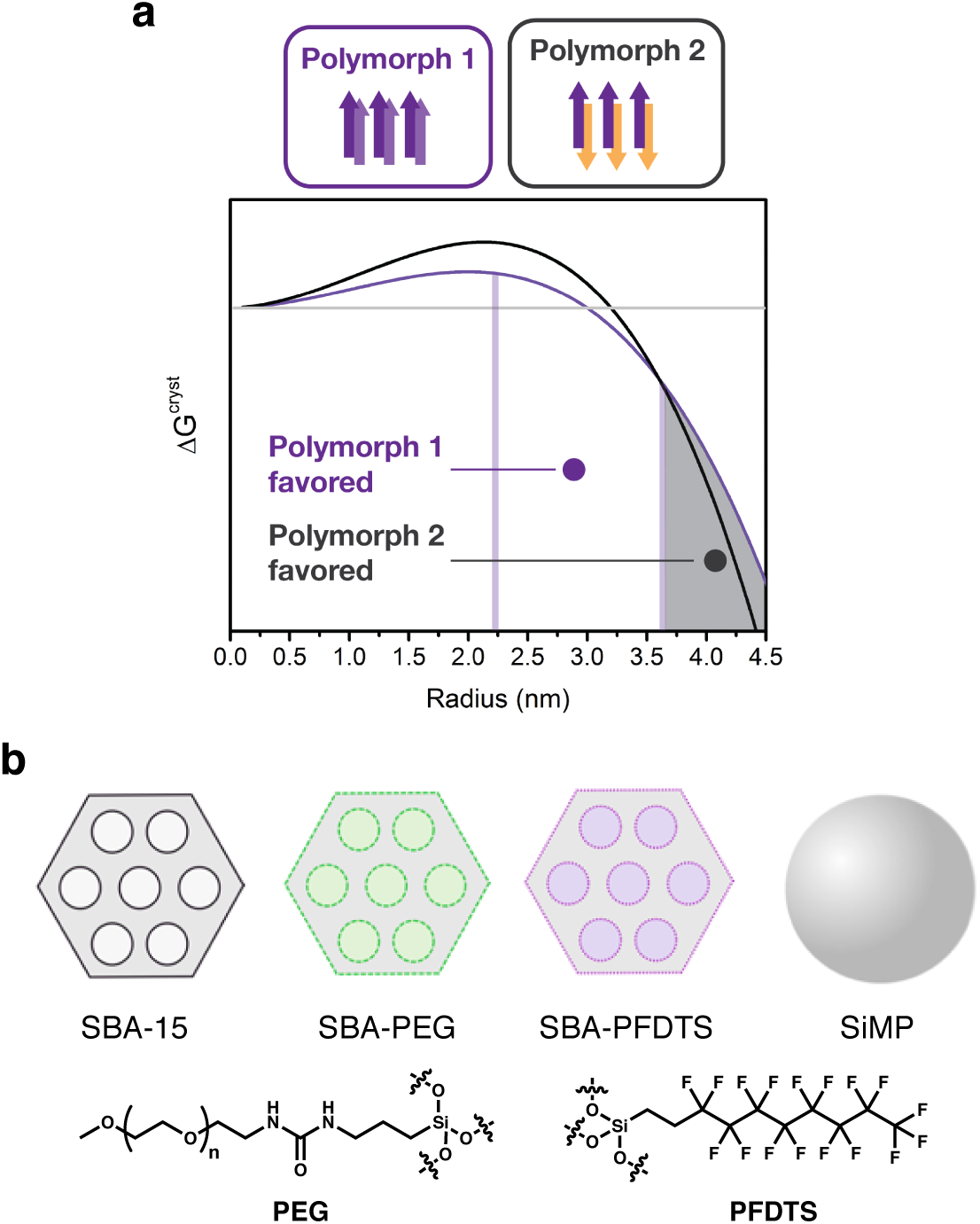
(a) Classical nucleation theory applied to the aggregation of Aβ. The critical nucleus is determined by the balance of bulk (volume) and interfacial forces. These forces ultimately determine the thermodynamic favorability of one polymorph over another. (b) The materials used in this study and the respective functional groups of SBA-PFDTS and SBA-PEG.

Here, we demonstrate that Aβ_1-40_ incubated with surface-functionalized mesoporous silicas (SBA-15) produces distinct amyloid fibril polymorph populations that depend on the silica surface chemistry. We characterized the interaction between the silicas and Aβ_1-40_ using a variety of techniques including fluorescence spectroscopy, IR spectroscopy, and atomic force microscopy. Subsequently, we used negative stain EM to identify different fibril polymorphs, which were defined by the number of twists, or cross-over features, per length of fibril. Using EM, we quantified relative fibril polymorph populations after Aβ_1-40_ was incubated with each material and found that the presence of specific fibril structures was highly dependent on silica surface chemistry and porosity. Specifically, SBA-15 and hydrophobically-functionalized SBA-15 accelerated Aβ_1-40_ aggregation kinetics and resulted in a higher population of fibrils with a shorter cross-over distance. Notably, Aβ_1-40_ fibrils with this feature are more structurally consistent with the structures of *ex vivo* fibrils. In contrast, buffer controls and hydrophilic-functionalized SBA-15 formed fibrils with either no cross-over or a 120 nm cross-over distance. Overall, our results suggest that synthetic materials can be leveraged for the *de novo* production of specific amyloid fibril populations and that EM is an ideal tool for characterizing heterogeneous populations of fibril polymorphs.

## Results

### Aβ Binding Affinity to Mesoporous Silicas

To probe the effect of exogenous materials on Aβ_1-40_ fibril polymorphism, we initially examined Santa Barbara Amorphous-15 (SBA-15), which is synthesized in a cooperative self-assembly process using a triblock copolymer consisting of ethylene oxide and propylene oxide units as a template followed by the addition of a silica source(45). The resulting mesoporous structure contains large cylindrical pores (4-30 nm), high surface area (>1000 m^2^ g^-1^), and tunable surface chemistry(46). These attributes are important because previous studies have shown that porosity and confinement influence nucleation and direct the resulting structure of small molecule crystals by changing the critical nucleation radius(47, 48). Moreover, SBA-15 has been shown to confine both myoglobin and lysozyme within its porous network, acting as an artificial chaperone that can affect protein structure(49, 50). In addition to leveraging its porosity and high surface area, many applications using SBA-15 require surface functionalization through post-synthetic grafting of organosilanes(51). Thus, we envisioned that the tunable surface chemistry of SBA-15 would allow us to tailor its interactions with Aβ_1-40_ and investigate how changes in the chemical and structural environment of the monomeric protein affect its aggregation into a threedimensional fibril structure.

Following SBA-15 synthesis, we used organosilane grafting to functionalize the material with a hydrophobic group, perfluorododecyltrichlorosilane (PFDTS), and hydrophilic group, polyethylene glycol (PEG) (Figure 1b). These functional groups were selected to evaluate the combined effects of surface charge, surface area, and hydrophobicity on Aβ_1-40_ aggregation kinetics and fibril structure. Finally, we also examined silica microspheres (SiMP, Cospheric SiO2MS-2.0.507μm) with a diameter of 500 nm as a control material to evaluate the effect of porosity on Aβ_1-40_ aggregation and fibril structure. Following synthesis and post-synthetic modifications, all materials were analyzed for surface area, surface charge, morphology, and surface chemistry (Table S1, Fig. S1-S3). Overall, using thermogravimetric analysis and NMR spectroscopy, we confirmed that surface grafting of the hydrophobic and hydrophilic groups was successful. Additionally, physisorption studies revealed that SBA-15 and SBA-PFDTS had the largest surface areas, while SBA-PEG had a reduced surface area available for adsorption.

Aβ binds to various proteins, metal ions, and cell receptors(52–54). Previous investigations have also examined the binding affinity of Aβ to exogenous materials, such as lipid vesicles(55). Because binding may influence effective protein concentration and structure, we first measured the binding affinity of Aβ_1-40_ to SBA-15 and its variants. Previous studies have shown that protein adsorption to SBA-15 can be modeled using a Langmuir adsorption model(49). Therefore, using a Bradford assay, we estimated the dissociation constant between monomeric Aβ_1-40_ and our SBA-15 variants. Both SBA-15 and SBA-PFDTS exhibited a relatively high affinity for Aβ_1-40_, with dissociation constants of 235 ± 27 μM and 112 ± 34 μM, respectively (Figure S4). The two-fold higher affinity for SBA-PFDTS is most likely a result of its hydrophobic nature and non-specific binding with the hydrophobic C-terminus of the Aβ_1-40_ monomer(56). These values are also consistent with previous studies that showed hydrophobic interactions dominated binding between short peptides and amyloids(57). In contrast, there was no measurable protein adsorption between Aβ_1-40_ and SBA-PEG and SiMP. The higher molecular weight of the PEG surface functionality reduced the surface area of SBA-PEG (Table S1) while also creating a hydrophilic surface less prone to interact with Aβ_1-40_. We measured a significantly lower surface area for the SiMP (6 m^2^ g^-1^), which potentially explains its lower affinity for Aβ_1-40_. Overall, these results confirm that surface functionality strongly influences the interaction of SBA-15 with Aβ_1-40_.

Next, we qualitatively validated our Aβ_1-40_ binding affinity measurements to the various materials through Western Blot (Figure S5). A mixture of Aβ_1-40_ monomer and pre-formed oligomers were incubated with each material for 15 minutes, followed by centrifugation, photocrosslinking, and SDS-PAGE. As expected, SBA-15 and SBA-PFDTS showed minimal Aβ_1-40_ monomer, oligomer, or fibril bands, indicating these materials effectively adsorbed and pulled down the monomeric protein and aggregates. In contrast, SBA-PEG displayed bands consistent with the Aβ_1-40_ control, suggesting minimal protein interaction with this material. Together, these results confirm that SBA-15 and SBA-PFDTS exhibit strong interactions with both Aβ_1-40_ monomers and aggregates.

### Kinetic effects of Mesoporous Silica on Aβ_1-40_ Aggregation

As mentioned above, a variety of environmental factors including exogenous materials, small molecules, and metal ions can influence the fibrillization kinetics of Aβ_1-40_. To determine the specific effect of our materials on the rate of Aβ_1-40_ aggregation, kinetic assays were conducted using thioflavin T (ThT) fluorescence, which is enhanced and red shifted in the presence of amyloid fibrils(58). Monomeric Aβ_1-40_ was incubated with each of the exogeneous materials in the presence of ThT and fluorescence spectra were collected for up to 20 hr (Figure S6). For the first 6-8 hr of incubation, no fluorescence was measured from any solution, indicating monomer association and dissociation in the absence of fibril formation (Figure 2)(59). Each solution then went through a rapid increase in ThT fluorescence, indicating rapid growth of mature fibrils due to fibril-catalyzed secondary nucleation(29). The length of the lag phase can be quantitatively estimated by modeling the fluorescence curves with a sigmoid function and evaluating the intercept of the tangent line at t1/2(60). Upon analysis of the lag time, we found that both porosity and surface chemistry influenced the aggregation kinetics of Aβ_1-40_. As seen in Figure 2, SBA-15 and SBA-PFDTS accelerated aggregation kinetics as indicated by a reduction in the length of the lag phase. Conversely, SiMP slightly slowed aggregation kinetics by extending the length of the lag phase, while SBA-PEG had no discernable effect, relative to the control, on the fibrillization rate. The rapid aggregation of Aβ_1-40_ in the presence of hydrophobic surfaces is consistent with previous studies, which found that these surfaces enhance *in vitro* and *in vivo* fibril self-assembly of Aβ_1-40_ and other amyloidogenic proteins(61, 62).

**Figure 2.**
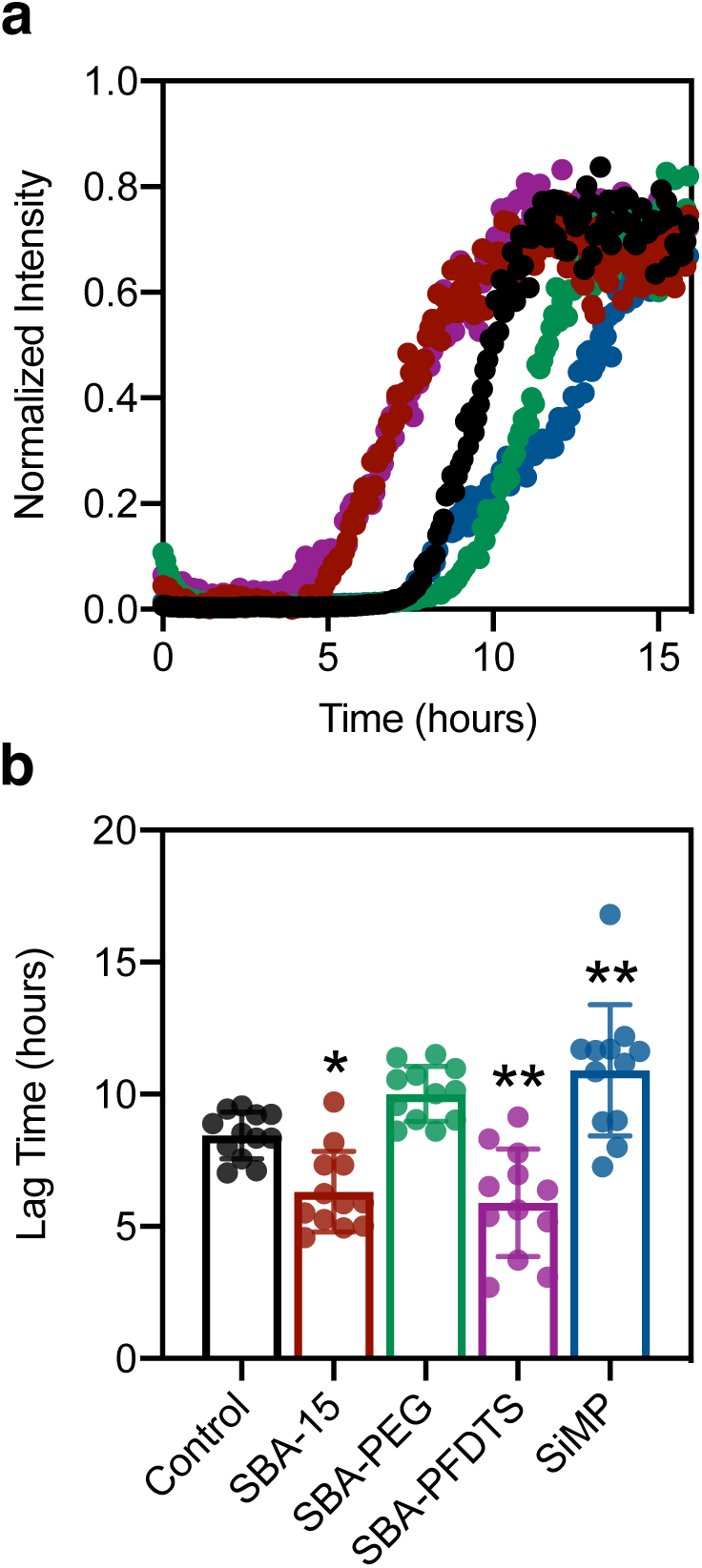
(a) Raw kinetic curves of the aggregation of 30 μM Aβ_1-40_ in 10 mM NaPi (black) at 37 °C and in the presence of 0.25 mg mL^-1^ SBA-15 (red), SBA-PEG (green), SBA-PFDTS (purple), and SiMP (blue). (b) Calculated lag time of aggregation for the aggregation. Fibrillization is accelerated in the presence of SBA and SBA-PFDTS. Error bars represent one standard deviation. *, p<.05. **, p<.01. ***, p<.001. ****, p<.0001 using a one-way ANOVA. Replicate kinetic data is presented in Figure S6.

### Preliminary Structural Effects of Mesoporous Silicas on Aβ_1-40_ Fibrils

Next, we investigated the effect of the silica materials on the structure of Aβ_1-40_ fibrils. First, infrared spectroscopy was used to determine if the β-sheets within the cross-β motif of the fibril were parallel or anti-parallel. Previous studies have shown that Aβ fibrils are largely comprised of in-register parallel sheets(63). However, anti-parallel sheets have been found in fibrils formed by smaller Aβ fragments and in amyloid-like crystals(63, 64). Both parallel and anti-parallel sheets have shown cytotoxic behavior, but the different arrangement of peptides into sheets could indicate different mechanisms of aggregation and self-assembly(63). Our IR measurements of the fibrils formed in the presence of the various materials led to a single peak at 1650 cm^-1^ (Figure S7), suggesting that parallel β-sheets were formed under all conditions tested.

To further probe the structural effects of mesoporous silicas on Aβ_1-40_, atomic force microscopy (AFM) was used to image the aggregates at various times in the fibrillization process. Relative to fluorescence measurements, AFM offers the advantage of being able to detect oligomers and smaller aggregates on the nanoscale that cannot be detected by other spectroscopic and microscopic methods(65). In order to examine the various aggregates using AFM, Aβ_1-40_ was incubated with SBA-15, SBA-PFDTS, SBA-PEG, and SiMP. Aliquots of the supernatant were taken at 2, 6, 8, and 24 hr to capture a variety of states of the aggregation process, and are shown in Figure 3. The SBA-15 and SBA-PFDTS trials both exhibited fibril formation at 8 hrs when compared to the control, indicating an acceleration in the kinetics. Conversely, SiMP inhibited fibril formation, with only small aggregates detected by AFM after 24 hours of observation (Figure S8). Ultimately, using AFM, we validated the kinetic trends observed in our ThT fluorescence experiments and confirmed that the Aβ_1-40_ behaves differently in the presence of the silicas with varying surface chemistry.

**Figure 3.**
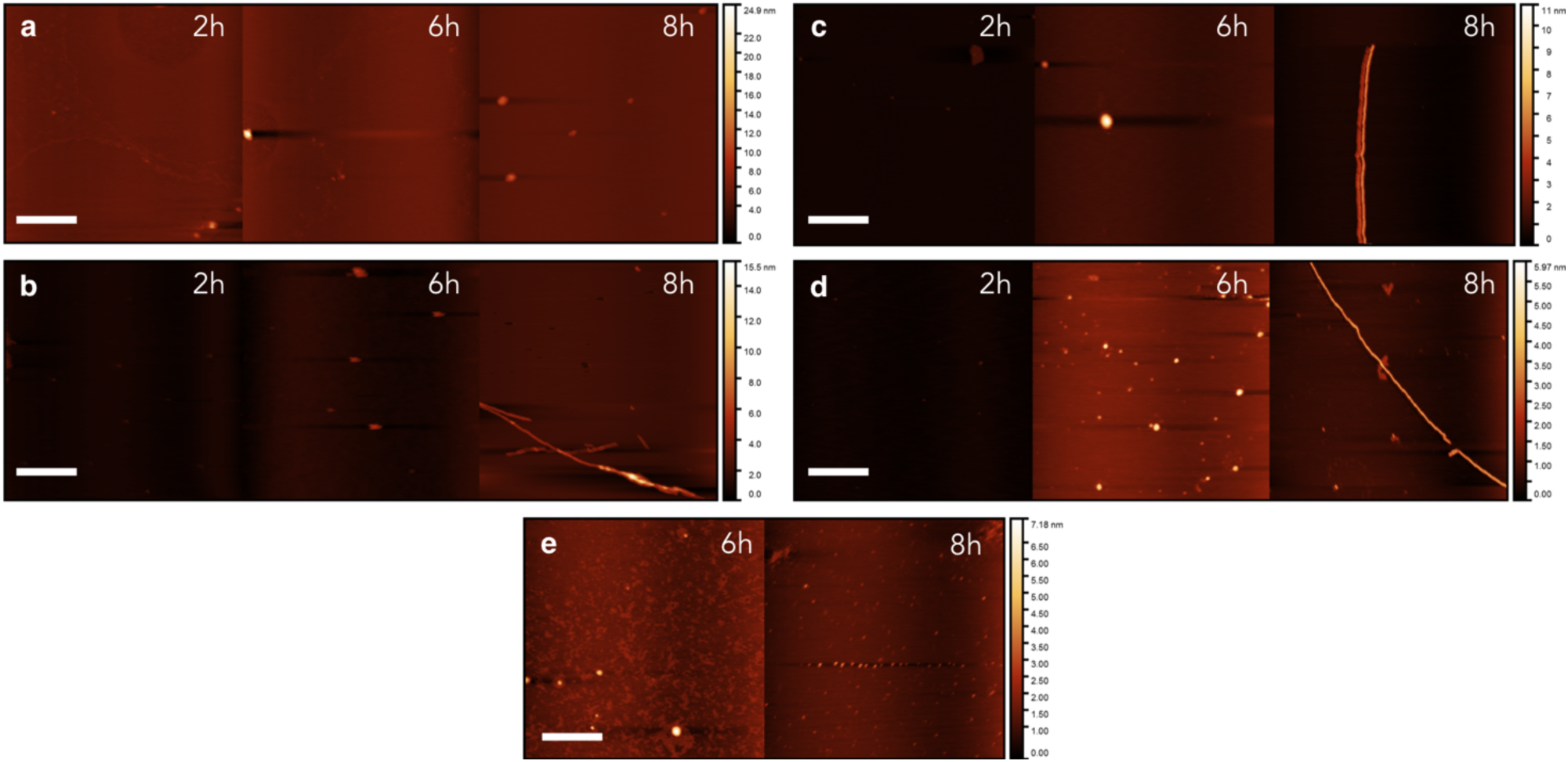
AFM Images of 30 μM Aβ1.40 after incubation in (a) 10 mM NaPi and 0.25 mg mL^-1^ of (b) SBA-15, (c) SBA-PEG, (d) SBA-PFDTS, and (e) SiMP at 37 °C after 2, 6, and 8 hr. Note that no oligomers were detected for SiMP at 2 hours. Scale bars are 500 nm. Images collected after 24 hr can be found in Supporting Information.

### Effect of Mesoporous Silicas on Fibril Polymorph Distribution

While IR and AFM provide basic insight into the structure and growth dynamics of amyloid fibrils(66), transmission electron microscopy can reveal the morphology and helical pitch of fibril structures. Additionally, EM provides an advantage over other structural methods by imaging a large field of individual fibrils in a single image that do not need to be purified to homogeneity. For example, previous EM studies revealed that Aβ_1-40_ fibrils take on various structures and the distribution of these polymorphs can change depending on the fibrillization conditions(23, 67, 68). Thus, we used negative stain EM to rapidly discern and quantify specific Aβ_1-40_ polymorphs formed in the presence of each exogenous material. We chose to use negative stain EM as samples were easily prepared and it offers a higher signal-to-noise ratio relative to cryo-EM.

To determine the effect of our materials on fibril polymorphism, Aβ_1-40_ monomer was incubated with each material for 24 hr. Each fibrillization experiment was conducted in triplicate to determine the reproducibility of morphological distributions formed in the presence of each exogenous material. Following incubation, fibril samples were collected by centrifugation, drop casted onto EM grids, and stained with uranyl acetate, U(OAc)_2_. Over 800 micrographs were collected from all experimental conditions, and *eman2helixboxer* from the EMAN2 image processing package (69) was used to extract fibril segments as individual particles with a box size of 123.84 x 123.84 nm. We chose this box size since the longest observed helical cross-over distance of the fibrils formed across all conditions was ~120 nm. Non-overlapping particles were selected from micrographs in order to accurately measure the population frequency of fibrils that adopted a specific cross-over distance. We then performed reference-free 2D classification using RELION(70) to investigate the structural similarity of Aβ_1-40_ morphologies observed across each condition (Figure 4). The 2D class averages confirmed the internal consistency of structure within each fibril and external consistency between fibrils formed under different conditions (Figure S9). Across the entire sample set, fibrils with no cross-over or cross-over distances of 120 nm, 60 nm, and 40 nm were observed for Aβ_1-40_. To the best of our knowledge, there is no standard nomenclature for different Aβ_1-40_ fibril polymorphs; therefore, we labeled the observed fibrils as δ, ε, λ, and μ fibrils, respectively. Fibrils that markedly changed helical cross-over distance along different parts of its length were very rarely observed (ca. 0.1%). The observed structures derived from class averaging agree with previous studies investigating the structural polymorphism of Aβ_1-40_ fibrils(23).

**Figure 4.**
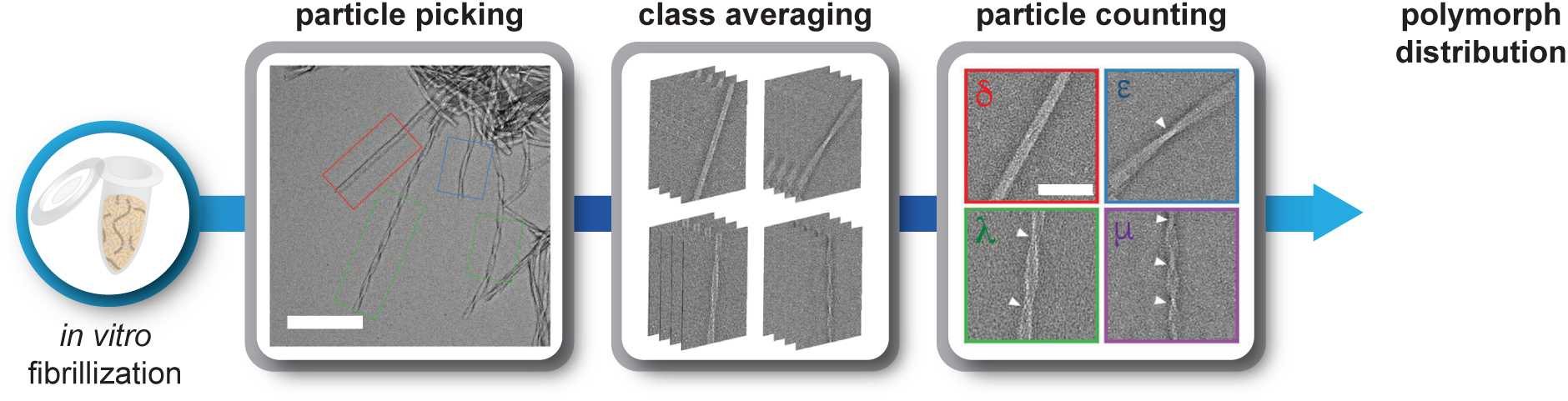
Workflow of the acquisition of fibril polymorph distributions. Aβ_1-40_ was incubated with each material for 24 hours at 37 °C. Fibrils were drop casted on TEM grids and stained for imaging. Particles displaying various morphologies were picked with a box size of 123.84 nm. The various morphologies were split into 4 classes based on a cross-over distance of 0 nm (δ), 40 nm (μ), 60 nm (λ), and 120 nm (ε). Particles were then counted and used to calculate the overall polymorph distribution in each condition.

In the Aβ control and SBA-PEG conditions, we observed primarily δ fibrils (no cross-over), ε fibrils (120 nm cross-over distance), and a small population of λ fibrils (60 nm cross-over distance). Specifically, the Aβ control produced 20.5 ± 9.2% δ fibrils, 72.5 ± 13.6% ε fibrils, and 7.1 ± 6.3% λ fibrils. This was comparable to the SBA-PEG, which produced 24.1 ± 17.5% δ fibrils, 71.6 ± 22.2% ε fibrils, and 4.3 ± 4.9% λ fibrils. Notably, these values are also consistent with previous literature reports describing the structure of fibrils formed under comparable conditions(68). With SBA-15, we observed a decrease in the percentage of ε fibrils at 32.1 ± 9.1%, which was offset by an increase of δ and λ fibrils at 38.9 ± 16.6% and 23.5 ± 13.6%, respectively, as well the appearance of a small population of μ fibrils (40 nm cross-over distance), at 5.5 ± 7.9%. SBA-PFDTS produced a distribution of fibrils that was significantly different from all other material conditions. Specifically, there was a large increase in the percentage of λ fibrils compared to the rest of the conditions, where 49.0% ± 10.3% λ fibrils were present. The polymorph distributions displayed in both the SBA-15 and SBA-PFDTS conditions, with an overall decrease in cross-over distance, were the most consistent to that of fibrils obtained *ex vivo*, albeit with the opposite helicity(22). Finally, the SiMP produced a population similar to the control, with a small population of μ fibrils observed, indicating the potential importance of porosity and surface area in the nucleation process.

Because both SBA-15 and SBA-PFDTS accelerated the kinetics of amyloid formation, we next examined whether their unique fibril population distributions were the result of a purely kinetic effect. For example, we asked if alternative incubation conditions that also accelerate fibril formation result in a similar polymorph distribution to SBA-PFDTS. To test this, Aβ_1-40_ monomer was incubated with sodium dodecyl sulfate (SDS), which is known to accelerate the aggregation of Aβ(71). As expected, SDS accelerated fibrillization kinetics with a comparable lag time to SBA-PFDTS (Figure S11). However, as seen in Figure 5, SDS produced a population of fibrils distinct from that of SBA-PFDTS, indicating that the observed fibril populations are the result of material effects beyond the rate of aggregation.

**Figure 5.**
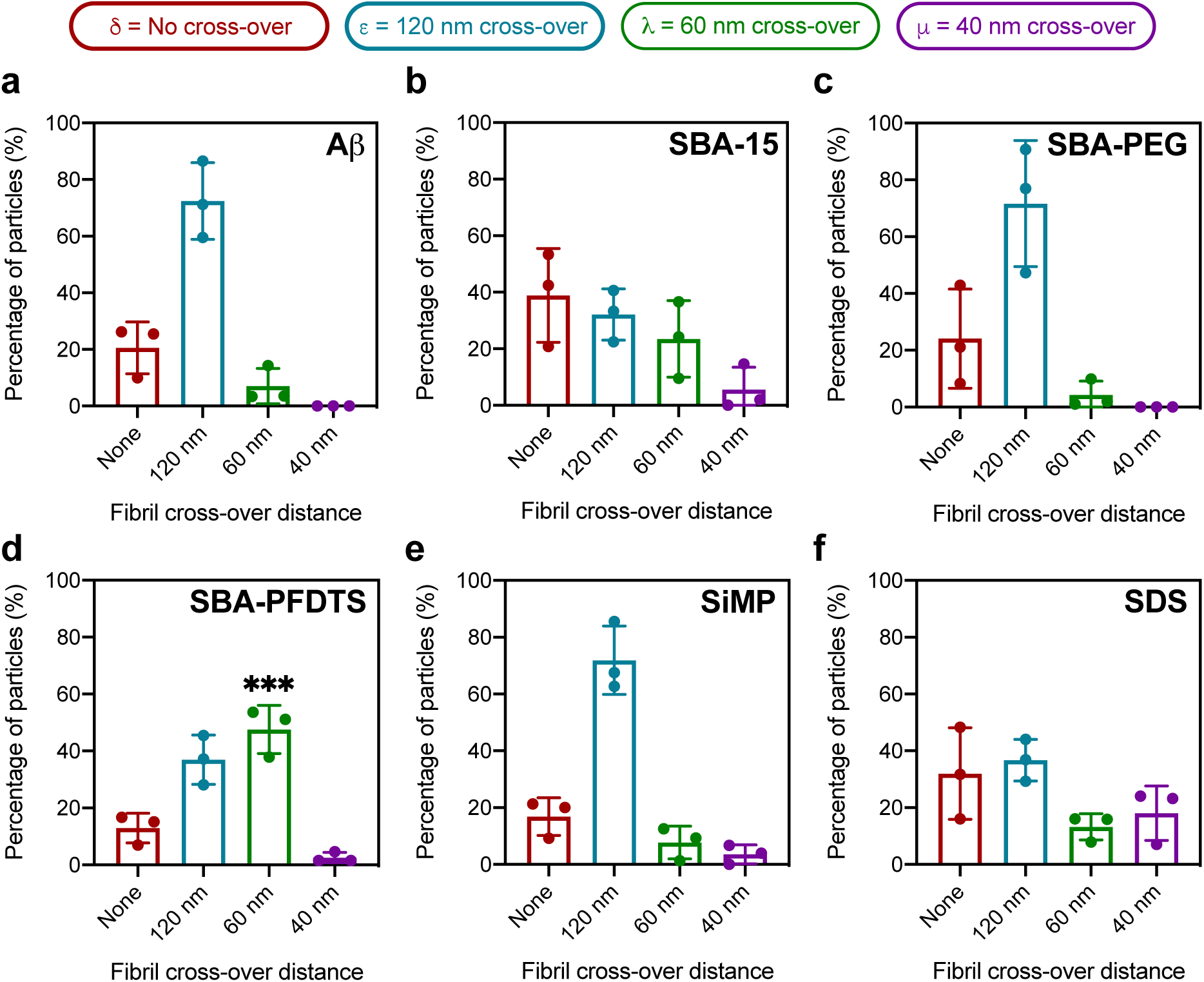
Distributions of cross-over distance in Aβ1.40 fibrils after a 24 hr incubation with (a) 10 mM NaPi buffer and 0.25 mg mL^-1^ of (b) SBA-15, (c) SBA-PEG, (d) SBA-PFDTS, (e) and SiMP. To test if these distributions were truly the result of a material effect rather than accelerated kinetics, distributions were also measured in the presence of (f) 100 μM SDS. Population fractions of each polymorph were averaged over 3 biological replicates (independent fibrillization experiments), and error bars represent 1 standard deviation. SBA-PFDTS produced a λ population significantly different from the control. ***, p = 0.0002 using one-way ANOVA.

## Discussion

We demonstrated that changing the surface area and functionalization of mesoporous silicas impacted the aggregation kinetics and structure of Aβ_1-40_ fibrils. In the case of ThT assays, we found that the addition of materials with differing surface chemistries altered the aggregation kinetics of fibril formation. Accelerated kinetics and a shorter lag phase were observed when Aβ_1-40_ was incubated with SBA-15 and SBA-PFDTS. In contrast, no significant changes in aggregation kinetics or the length of the lag phase were observed when Aβ_1-40_ was incubated with SBA-PEG. We attribute these accelerated kinetics in SBA-15 and SBA-PFDTS to the high surface area of SBA-15 (886 m^2^/g) and the additional hydrophobicity of SBA-PFDTS(61, 62). In contrast, the relative hydrophilicity of the SBA-PEG mitigated any acceleration effects, yielding a lag time similar to that of the Aβ_1-40_ control. For comparison, we also tested the effect of SiMPs, which have a comparable particle diameter to the SBA-15 variants but a lower surface area (6 m^2^/g). We hypothesize that the lower surface area of the SiMPs is responsible for their negligible effect on amyloid kinetics. To confirm these results, we also conducted AFM measurements to visualize Aβ_1-40_ oligomers and fibrils across a 24 hr period. The AFM results were consistent with the aggregation kinetics observed in the ThT assays (Figure 3). Overall, our kinetic and biophysical studies agree with previous reports examining the effect of exogenous materials and hydrophobic interfaces on the aggregation kinetics of amyloid formation, while providing additional insight into the effect of porosity, surface area, and surface chemistry on amyloid fibrillization.

Our main goal was to investigate the effect of mesoporous silicas and influencing nucleation on Aβ_1-40_ fibril polymorphism. Similar to other amyloidogenic proteins, Aβ_1-40_ fibrils adopt a range of structures, even under relatively simple incubation conditions(10, 23)(21)(72). Structural polymorphism between fibrils can include changes in protein orientation, cross-over distance, and nucleation kinetics(73). However, the most common way to differentiate structure is by helical pitch (cross-over distance). Several polymorphs for amyloidogenic proteins formed under *in vitro* and *ex vivo* conditions have been recently characterized at atomic resolution with cryo-EM(74–76). As observed in recent *ex vivo* studies(22), it is unclear how representative amyloid fibrils prepared *in vitro* are of structures found in patients thus far.

Nucleation theory and amyloid specific work have suggested that influencing the nucleation environment can potentially control the distribution of fibril structures(67, 77). To test this hypothesis, we analyzed the population distribution of different fibril polymorphs formed in the presence of functionalized mesoporous silicas using negative stain EM and 2D classification. The use of negative stain EM, instead of cryo-EM, for quantifying fibril populations offered a relatively high-throughput method for imaging and visualizing structural differences across incubation conditions, which allowed us to test a variety of nucleation environments in a statistically significant manner. To rapidly classify different polymorphs, we sorted fibril particles based on different cross-over distances, since this feature was most distinct when differentiating polymorphs during the class-averaging process. Initial class-averages were further refined through subclassification. Through this analysis, we observed significant differences in the distribution of amyloid fibril polymorphs for each incubation condition and that changes to the exogenous material’s porosity and surface functionalization influenced the frequency of one fibril structure compared to another. Overall, we observed fibrils with no cross-over (δ-Aβ_1-40_) and cross-over distances of 120 nm (ε-Aβ_1-40_), 60 nm (λ-Aβ_1-40_), and 40 nm (μ-Aβ_1-40_).

As shown in Figure 5, Aβ_1-40_ incubated under control conditions and in the presence of SBA-PEG or SiMP primarily produced δ and ε fibrils. Previous reports using EM and SSNMR spectroscopy identified amyloid fibrils with similar structures when Aβ_1-40_ was incubated in a variety of simple buffers. The inclusion of different salt ions also produced similar distributions of δ and ε fibrils(78). The δ fibrils are dimeric in each fibril repeat(39), while the ε fibrils can contain two to four peptides in each repeat(17)(79). Additionally, the reported atomic resolution structures show that δ fibrils have parallel, in-register β-sheets, with the majority of hydrophobic amino acid sidechains residing in the interior of the fibril and the majority of the polar amino acid residues residing on the exterior. Polar zipper interactions may also occur with residue Q15 fitting into the cavity created by Q37 and Q38. For the ε fibrils, as shown in Figure 6, a trimeric structure has been reported in which each cross-over represents a 60° rotation. In this structure, the peptide conformation is similar to that of the δ fibrils, where the majority of the hydrophobic amino acid sidechains reside in the core(13). Overall, the observed dominance of δ and ε fibril structures in our control, SBA-PEG, and SiMP samples is consistent with previous literature reports and validates our methodology for fibril classification.

**Figure 6.**
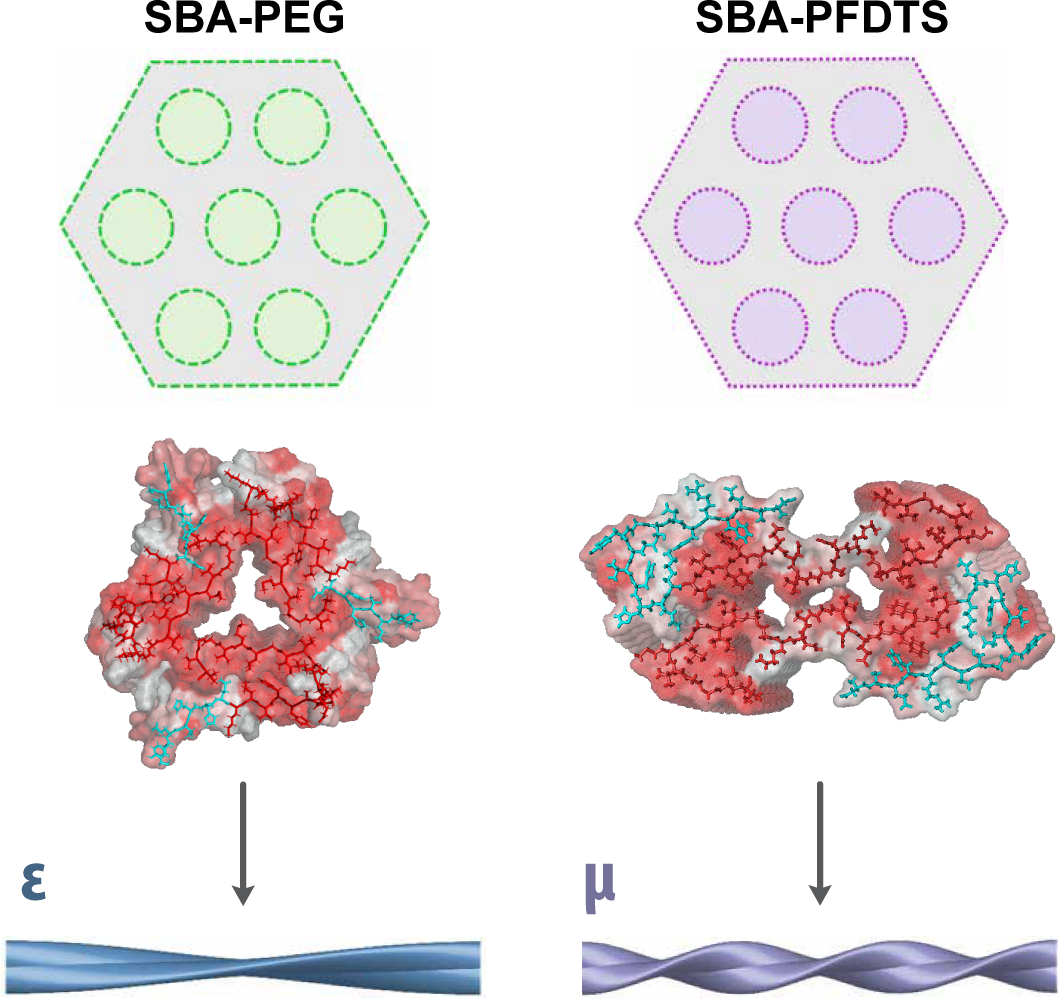
Representative atomic structures of the fibril morphologies observed for SBA-PEG and SBA-PFDTS. Hydrophobic residues are shown in red using *color_h* in PyMol(83). ε-fibrils with a cross-over distance of 120 nm (PDB: 2LMQ) were primarily observed in the control and in the presence of SBA-PEG. Populations with smaller cross-over distances, λ and μ, were observed in the presence of SBA-PFDTS, for which an *ex vivo* fibril with a similar morphology has been observed (PDB: 6SHS). In the polymorph with a smaller cross-over distance (μ), hydrophobic residues were exposed to the solvent, suggesting that the hydrophobic environment of SBA-PFDTS could be influencing the nucleation.

Consistent with our hypothesis that the nucleation environment can influence fibril structural polymorphism, we observed that SBA-15 and SBA-PFDTS had a unique effect on the distribution of fibril polymorphs. SBA-15 showed a decrease in ε fibrils coupled with an increase of both δ and λ fibrils, as well as the appearance of a population of μ fibrils. SBA-PFDTS exhibited a statistically significant increase in the proportion of λ fibrils compared to all other materials tested. Fibrils with a morphology (cross-over distance) similar to our observed μ fibrils were recently identified in an *ex vivo* study involving AD patients, albeit with an opposite helicity to those found in our experiments(22). These μ fibrils have dimeric fibril repeats, a C-shaped peptide fold, and a central peptide domain that is buried in the fibril core, which interacts between the two peptide stacks at residues 24-26. In contrast to δ and ε fibrils, the majority of charged amino acid side chains in the μ fibril structure are solvent exposed, and Q11 and K16 are oriented in a manner that can form salt-bridges with one another. Small populations of λ fibrils have been observed in the previous studies but the atomic structure has yet to be reported(79).

Previous molecular dynamics (MD) simulations of fragments of amyloidogenic protein sequences (Sup35) have suggested that changes in fibril twisting (cross-over distance) may be due to different charge interactions at the protein’s N- and C-termini(80, 81). Specifically, with a negative charge at the C-terminus, model proteins adopted the more energetically favorable twisting conformation in MD simulations compared to the fibrils with no cross-over distance, which were less energetically favored(81). These simulation results may explain why the accessible negative surface charge of SBA-15 favors the formation of λ and μ fibrils compared to the ε fibrils that were highly represented in the control, SBA-PEG, and SiMP samples. Similarly, the larger population of λ and μ fibrils observed in the presence of SBA-PFDTS indicates that hydrophobicity plays a significant role in determining fibril structure and we speculate that hydrophobic environments may generally favor these structures. Analyzing the currently available PDB structures (2LMQ and 6SHS) corresponding to the ε fibrils and μ fibrils revealed that the hydrophobic C-termini in the μ fibril structure are more solvent-exposed than the ε fibrils, as shown in Figure 6(17, 22). Overall, this suggests that the exposure of different regions of Aβ_1-40_ to the functionalized materials during aggregation determines which polymorph is favored.

It is notable that while SDS accelerated the aggregation kinetics of Aβ_1-40_ and exhibited a moderate increase in the proportion of λ fibrils, the distribution was not as prominent as SBA-PFDTS. This suggests that the increased fraction of λ fibrils observed in the presence of SBA-PFDTS is not solely due to accelerated aggregation, and that the surface area and chemistry of the materials may be the driving mechanism for this morphology. Finally, we hypothesize that the negative charge introduced by the surfactant may have a similar an effect on Aβ_1-40_ as SBA-15, potentially yielding the population of μ fibrils that was not observed in the control and SBA-PEG.

Overall, our results show that mesoporous silicas influence the aggregation kinetics of Aβ_1-40_ and the distribution of observed fibril morphologies. By tuning the material properties of mesoporous silicas, we highlighted the potential of exploiting nucleation theory in conjunction with exogenous materials for biasing amyloid formation toward specific fibril polymorph distributions. Future refinement of our methodology could facilitate *in vitro* access to fibril structures more representative of neurodegenerative disease isolates or to specific fibril polymorphs with unique structures and pathologies.

## Materials and Methods

### SBA-15 Synthesis

4 g of Pluronic 123 (Mn-5,800, Sigma-Aldrich #435465) was mixed with 30 mL H2O and 70 mL of 2M HCl for 1 hr at room temperature. The mixture was then brought to 40 °C and 9 g of tetraethyl orthosilicate (98%, ACROS Organics #1577812500) was added. The reaction mixture was stirred for 24 hr, and then vacuum filtrated. The resulting product was aged in an oven at 100 °C for an additional 24 hr. To remove residual surfactant from the pores, samples were calcined at 600 °C for 8 hr.

### SBA-15 Functionalization

500 mg of SBA-15 was mixed with 110 mg of 1H,1H,2H,2H-Perfluorododecyltrichlorosilane (PFDTS) (Sigma Aldrich #729965) or 160 mg of mPEG-Silane (Mn-1,000, Laysan Bio Inc.) in 35 mL dry toluene in a nitrogen atmosphere. The reaction was then carried out under an argon atmosphere at 110 °C under reflux for 20 hr. The resulting product was then vacuum filtrated and dried under high vacuum on a Schlenk line.

### Surface Area Measurements

Using an ASAP 2020 Physisorption (Micromeritics), BET surface area was measured for the various mesoporous silica, using He gas for free space measurements and N2 gas for adsorption. Samples were activated at 120 °C under high vacuum (10 μbar) for 24 hr prior to the BET measurement.

### Zeta Potential Measurements

Using a Dynamic Light Scattering Zetasizer (Malvern 2010), zeta potential was measured using a suspended mixture of mesoporous silica (0.25 mg/mL) in 10 mM sodium phosphate (NaPi) buffer at 25 °C. Zeta potentials are represented as the average of triplicates, each with 20 minimum scans.

### NMR of Functionalized SBA

Functionalization of the SBA-15 was confirmed using an AVANCE III 400 solid state NMR, running C^13^ NMR at a spin of 8 kHz. Resulting spectra were evaluated using MesterNova.

### Preparation of Aβ_1-40_

5 mg of Aβ_1-40_ (BACHEM, H-1994) was dissolved in 500 μL of hexafluoro-2-propanol (HFIP) and shaken at 300 RPM for 1 hr. An additional 500 μL of HFIP was added and 40 μL aliquots were distributed into microcentrifuge tubes. The HFIP was allowed to evaporate overnight and subsequently dried using a vacuum centrifuge. The samples were stored at −20 °C.

### Kinetic Assays of Aβ_1-40_ Aggregation

An aliquot of monomer (230 μg) was thawed and then dissolved in 20 μL of DMSO and 880 μL of 10 mM sodium phosphate (NaPi) buffer to produce a 60 μM solution of Aβ_1-40_. Aliquots of dissolved monomer (100 μL) were then pipetted into a nonbinding, coated 96 well plate (Corning 3991). Each well was then diluted with 100 μL of a suspension containing 0.5 mg mL^-1^ of exogenous material and 200 μM of Thioflavin T (ThT) in 10mM NaPi. The final volume of each well was 200 μL with a concentration of 30 μM Aβ_1-40_, 100 μM ThT, and 0.25 mg mL^-1^ of exogenous material in 10 mM NaPi. The 96 well plate was sealed with spectroscopy grade tape (Thermo Scientific #235307) and placed in a plate reader (Clariostar, BMG Labtech) at 37 °C. Assays were conducted with shaking at 300 RPM. ThT fluorescence intensity was measured with an excitation of 440 nm and an emission of 480 nm through the top of the plate. Measurements were taken every 300 seconds for 20 hours. Each sample was run in triplicate with blanks (without Aβ_1-40_) to account for ThT interactions with the exogenous material.

### Lag Time Calculations

The lag time for each kinetic assay was calculated by fitting a sigmoidal curve to the fluorescence data of the aggregation (Eq. 1),

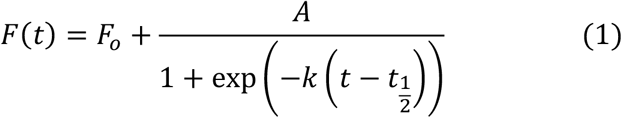

where *A* is the amplitude of the largest fluorescence intensity, *F_o_* is the background fluorescence, and *t_1/2_* is the time at which the intensity is ½ of the max value.^8^ The lag time, *t_lag_*, is defined as the intercept of the tangent line at *t_1/2_* with slope *k* (Eq. 2):

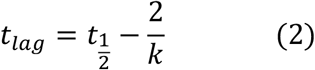

### Binding Affinity Bradford Assay

To evaluate the binding affinity of Aβ_1-40_ for the surface and pores of the nanoparticles, a 200 μL solution of 30 μM Aβ was mixed with the various materials ranging in concentration from 0.1 to 15 mg mL^-1^ in a microcentrifuge tube. These suspensions were incubated for 15 min at room temperature, shaking at 300 RPM. The samples were subsequently centrifuged, and 20 μL of the supernatant was added to 180 μL of Coomassie Plus (Thermo Scientific #23238). Absorbance measurements were taken at 595 nm in triplicate, and the absorbance intensity was then compared to a calibration curve to determine the protein concentration of the supernatant. Association constants were calculated by modeling the absorption curve to a Langmuir isotherm.

### Binding Affinity Western Blot Assay

30 μM of Aβ_1-40_ monomer was incubated for 20 hr at 4 °C to form oligomers. The pre-formed oligomers were then incubated with each material (0.25 mg mL^-1^) for 15 min at room temperature in a 1.5 mL microcentrifuge tube, and the suspension was subsequently centrifuged. The supernatant was then collected and photo-crosslinked using the PICUP reaction(82). 18 μL of the supernatant were mixed with 1 μL of 3 mM tris(2,2’-bipyridyl)-dichlororuthenium(II) and 1 μL of 60 mM ammonium persulfate in a 0.2 mL PCR tube. The mixture was irradiated with an LED light for 10 s and quenched with 20 μL of a 5% solution of β-mercaptoethanol. Samples were prepared for SDS page, by adding dye and then heating at 95 °C for 7 min. After SDS-PAGE, the gel was transferred to a 0.2 μm PVDF membrane and blocked in 4% milk. The membrane was then incubated with 6E10 (1:1000) and Goat antimouse HRP (1:2500).

### Atomic Force Microscopy

Samples were prepared by centrifuge-spinning the incubated Aβ from the different types of functionalized mesoporous silica, at 15,000 rpm for 2 min. After separating the Aβ-containing supernatant from the mesoporous silica pellet, a 100μL aliquot was deposited onto a piece of freshly-cleaved mica (Ted Pella, highest grade mica sheets) and incubated at room temperature for 1 min. The excess liquid was then removed from the edge of the mica with a Kimwipe and the surface was rinsed with a 100μL aliquot of Dulbecco’s phosphate buffered saline (Sigma) before it was dried under a stream of N2(g). An Asylum MFP-3D atomic force microscope was used to image dropcasted Aβ on freshly-cleaved mica. AFM cantilevers (MikroMasch) with a typical probe radius of 8 nm, 65 kHz resonance frequency, and 0.5 N m^-1^ force constant were used to analyze the surface under tapping mode, to minimize sample damage due to shear forces and tip-sample interactions. All images were processed using the Gwyddion SPM software package.

### Attenuated Total Reflectance Infrared Spectroscopy (ATR-IR)

Vibrational spectroscopy was collected with a Bruker Vertex 70 Fourier transform infrared (FTIR) spectrometer equipped with a VeeMAX II ATR accessory (Bruker) to illuminate the sample. The samples were centrifuged at 15,000 rpm for 2 min to separate the concentrate the Aβ suspension. Using a centrifugal evaporator to remove the liquid, the Aβ was then resuspended in 10 μL of 10 mM sodium phosphate buffer (NaPi) with a pH of 7.4 to a concentration of 2mM and deposited onto a ZnSe ATR crystal. After purging the sample chamber with N2(g) for 30 min, 500 scans of light were collected from each sample. A mercury cadmium telluride (MCT) detector was used to collect signals from 800-3000 cm^-1^. For each sample, a background sample of supernatant from an incubation using only mesoporous silica was prepared identically to the process described above in order to determine the absolute difference of absorption due to Aβ. Following this background subtraction, spectra were flattened using a rubber band correction baseline function in the instrument’s OPUS software.

### Transmission Electron Microscopy

Transmission electron microscopy was used to analyze the morphology of the mesoporous silica and the resulting Aβ_1-40_ fibrils. Samples were prepared on carbon coated grids (Electron Microscopy Sciences, CF400) that were glow discharged with an EmiTech K100x Coater. After allowing the amyloid samples to aggregate for 24 hr, 7 μL of the amyloid-mesoporous silica suspension was dropcast on the charged carbon side of the grid. After incubating for 1 min, the droplets were washed two times each with in 50 μL of 0.1 M and 0.01 M ammonium acetate. The droplet was then washed with 50 μL of 2% uranyl acetate. Following the washes, excess liquid was wicked away using filter paper. Samples were imaged using a JEOL 2010F TEM.

### RELION Class-Average of Fibril Structure

Negative stain images were acquired on a JEOL 2010F TEM operated at 200kV at a nominal magnification of X60K, which resulted in a pixel size 3.6 Å/pixel. The exposure for each image was 2 seconds, resulting in a total electron dose of 60-70 e^-^Å^-2^. Data was collected on a Gatan OneView camera with a defocus ranging from −0.5 μm to −4.0 μm.

### Image Processing

Fibril segments were picked using *eman2helixboxer* from the EMAN2 image processing suite, with a box size of 344 pixels and no overlap. This box size was selected based on the measurement of the longest helical cross-over distance observed. Fibril particles initially were manually picked using EMAN2 from 100 micrographs of each of six incubation conditions which were then extracted and segmented into particles with a box size of 1,238.4 Å using RELION. Approximately 520 particles were used to measure the distributions of fibrils with different helical cross-over distance across conditions. The particles were further refined by 2D classification into 3 classes using RELION with helical symmetry optimization enabled. Additional replicates were each comprised of 20 micrographs for each condition and processed using the same workflow described above.

## Acknowledgements

This work was supported in part by Welch Foundation Research Grants F-1929 (to BKK), F-1722 (to LJW), and F-1938 (to DWT), National Science Foundation CHE-1807215 (to LJW), Army Research Office Grant W911NF-15-1-0120 (to DWT), a Robert J. Kleberg, Jr. and Helen C. Kleberg Foundation Medical Research Award (to DWT). Partial support was provided by the National Science Foundation through the Center for Dynamics and Control of Materials: an NSF Materials Research Science and Engineering Center under DMR-1720595. D.W.T is a CPRIT Scholar supported by the Cancer Prevention and Research Institute of Texas (RR160088) and an Army Young Investigator supported by the Army Research Office (W911NF-19-1-0021). NMR spectra were collected on a Bruker Avance III HD 400 funded by the National Science Foundation (CHE 1626211). We thank Andres Sanchez-Paiva and Zhichao Chen for their experimental assistance. We would also like to thank Prof. Joan Brennecke for the use of her Thermogravimetric Analyzer. Additionally, we are grateful for the use of the facilities within the Texas Materials Institute and the technical expertise provided by Dr. Karalee Jarvis. Finally, we’d like to thank the core microscopy laboratory of the Institute for Cellular and Molecular Biology at the University of Texas at Austin.

