## Supplementary Info for "Functionalized Mesoporous Silicas Direct Structural Polymorphism of Amyloid-β Fibrils"

PNAS
www.pnas.org

Supplementary Information for

### Functionalized Mesoporous Silicas Direct Structural Polymorphism of Amyloid-β Fibrils

Michael J. Lucas[1†], Henry S. Pan[1†], Eric J. Verbeke[2†], Lauren J. Webb[3*], David W. Taylor[2,4-6*], and Benjamin K. Keitz[1*]

[1]McKetta Department of Chemical Engineering, University of Texas at Austin, Austin, TX, 78712
[2]Institute for Cellular and Molecular Biology, University of Texas at Austin, Austin, TX, 78712
[3]Department of Chemistry, University of Texas at Austin, Austin, TX, 78712
[4]Department of Molecular Biosciences, University of Texas at Austin, TX, 78712
[5]Center for Systems and Synthetic Biology, University of Texas at Austin, TX, 78712
[6]LIVESTRONG Cancer Institutes, Dell Medical School, Austin, TX, 78712

Lauren J. Webb, David W. Taylor, Benjamin K. Keitz

**This PDF file includes:**

Figures S1 to S11
Table S1

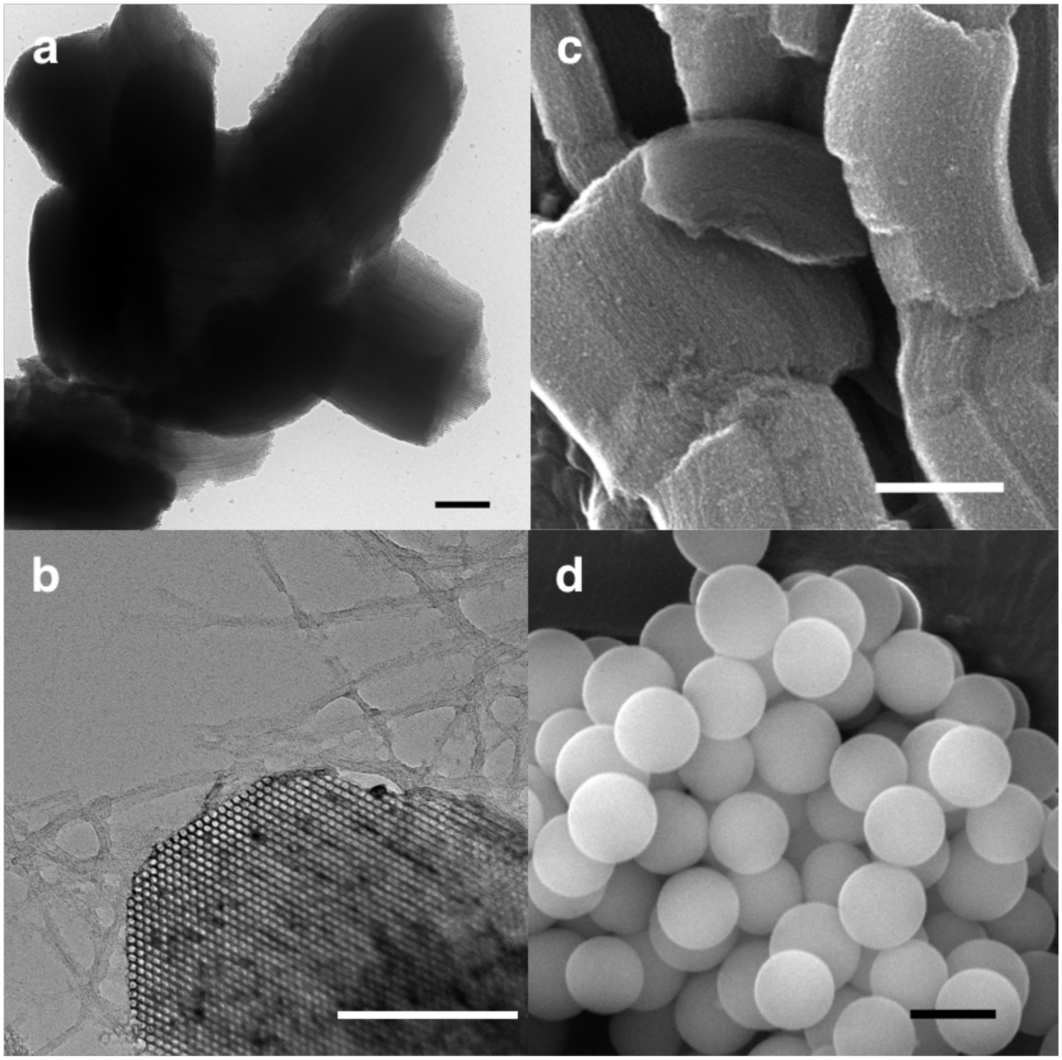


**Figure S1**. (a) TEM image of SBA-15. Scale bar is 200 nm. (b) TEM image of SBA-PEG axially with amyloid fibrils. Scale bar is 200 nm. (c) SEM image of SBA-15. Scale bar is 500 nm. (d) SEM image of silica microspheres (SiMP) with diameter of 500 nm. Scale bar is 500 nm.

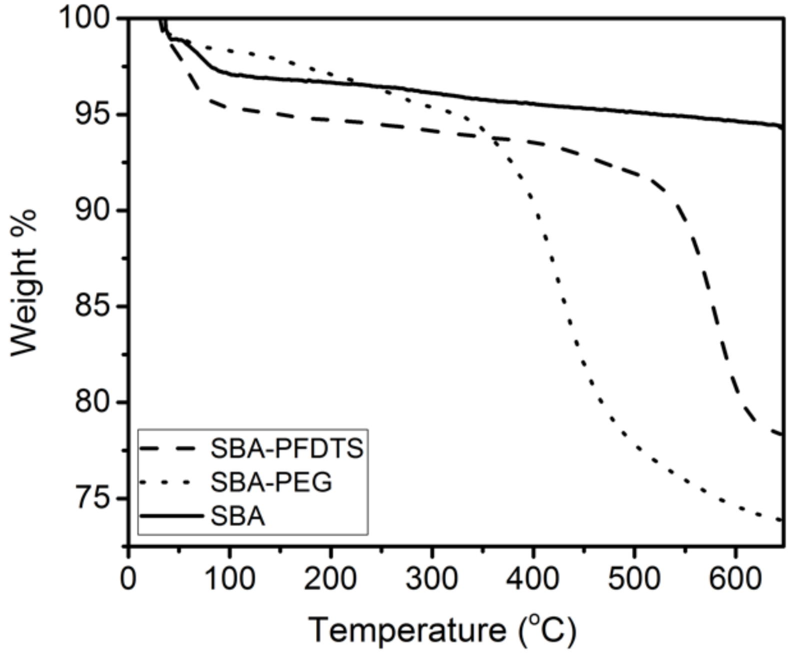


**Figure S2**. Thermogravimetric analysis (TGA) of functional groups, PFDTS (hydrophobic) and PEG (hydrophilic) attached to the surface of SBA-15.

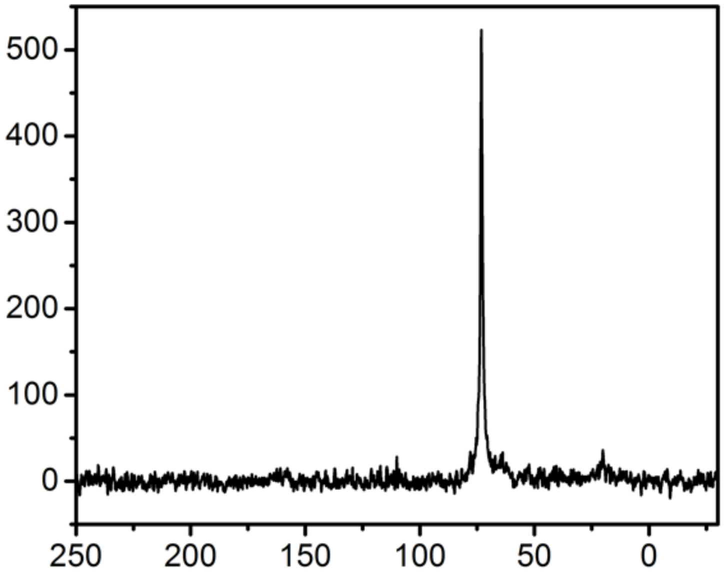


**Figure S3.** Solid State $C^{13}$ NMR spectrum of SBA-PEG.

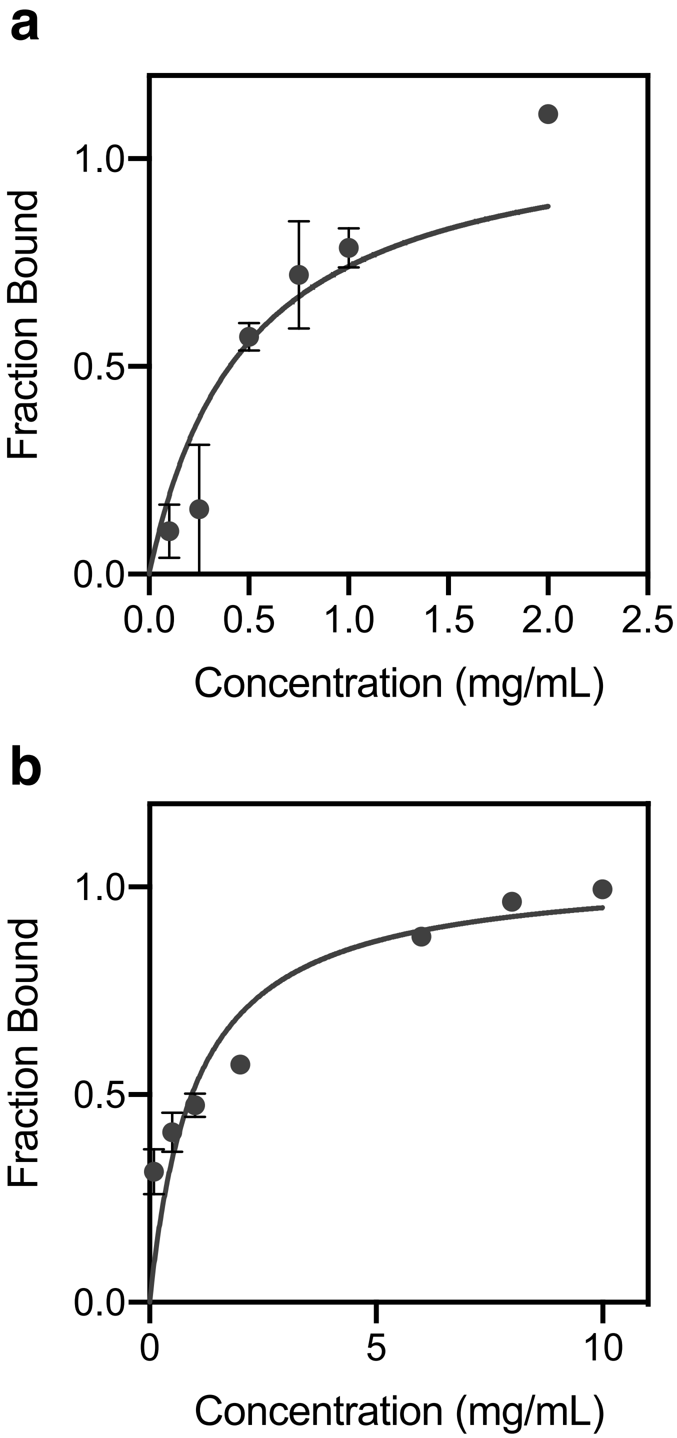


**Figure S4.** Langmuir Adsorption fits of A$\beta_{1\text{-}40}$ binding to (a) SBA-PFDTS and (b) SBA-15. Dissociation constants were 112 ± 34 μM and 235 ± 27 μM, respectively. Each concentration was run in triplicate. Error bars represent one standard deviation.

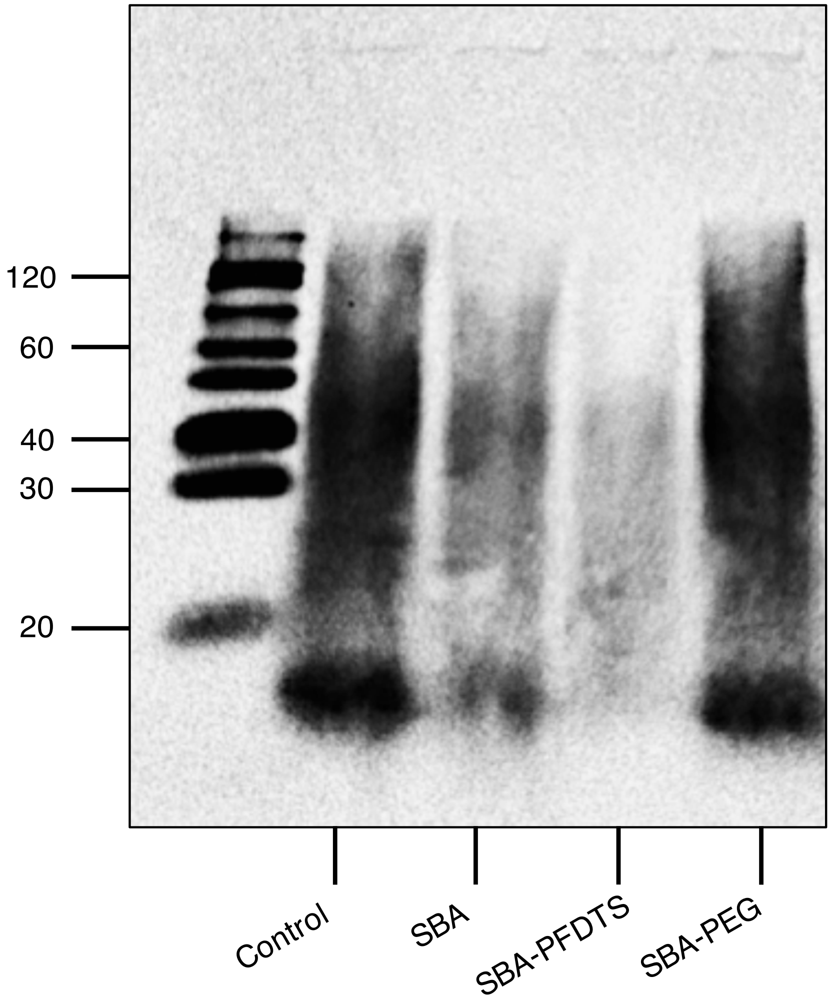


**Figure S5.** Western blot of pre-formed oligomers of A$\beta_{1-40}$ after a 15 minute incubation with the mesoporous silicas. SBA-PFDTS had the strongest binding affinity for A$\beta_{1-40}$, as evidenced by the decreased intensity of oligomer bands. SBA also showed an affinity for A$\beta_{1-40}$, while SBA-PEG showed no difference to the control, in agreement with the Bradford assays. Oligomers were crosslinked using the PICUP reaction. Primary antibody used was 6E10. Units for weights are shown in kDa.

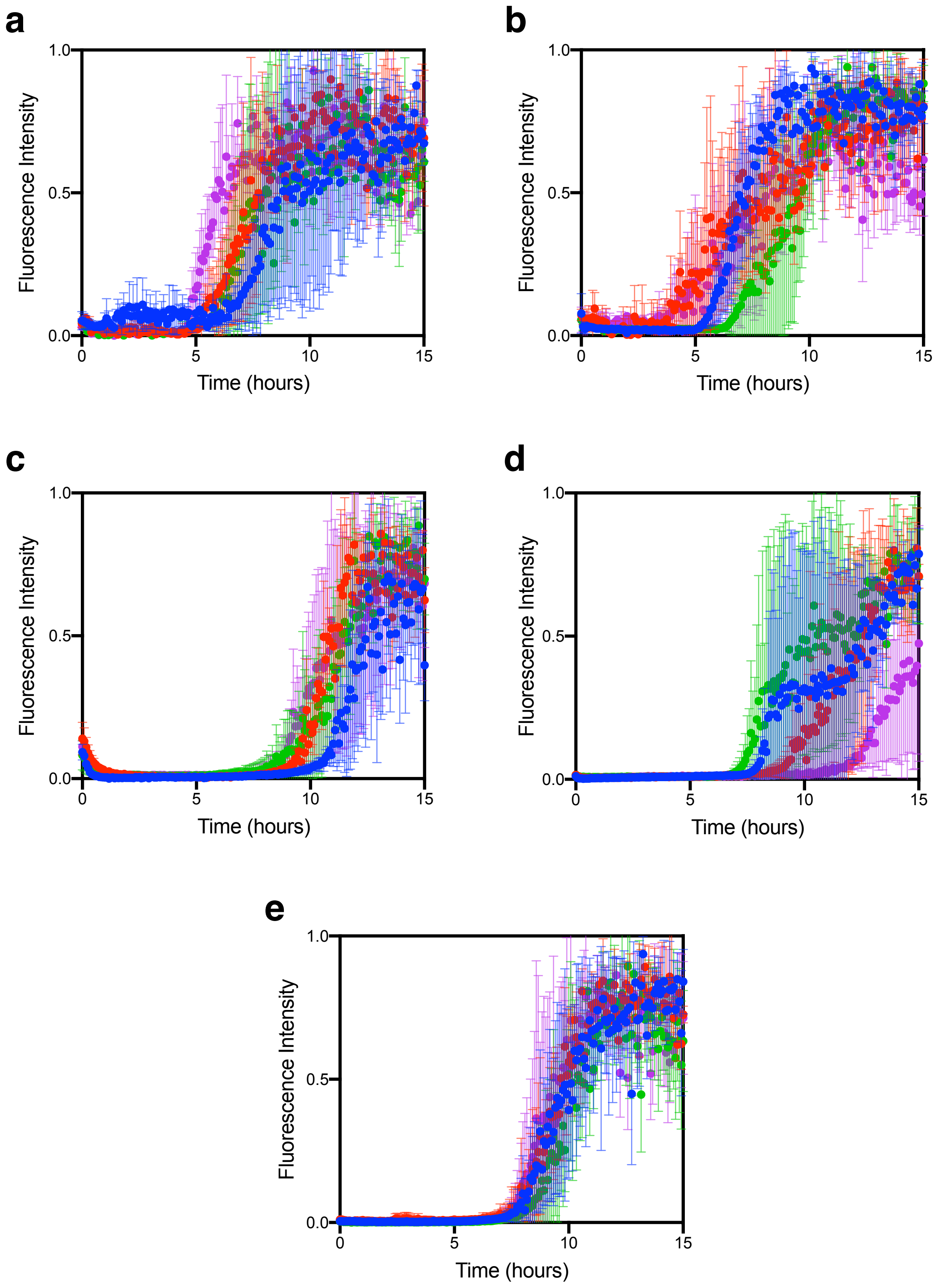


**Figure S6.** ThT Fluorescence curves of the fibrilization of A$\beta_{1\text{-}40}$ at 37 °C in the presence of 0.25 mg/mL of (a) SBA-15, (b) SBA-PFDTS, (c) SBA-PEG, (d) SiMP, and (e) 10mM NaPi buffer. For each material (plus control) 4 replicates were collected consisting of 3 kinetic runs each. Each color represents the mean ± SD of one experiment (3 kinetic runs).

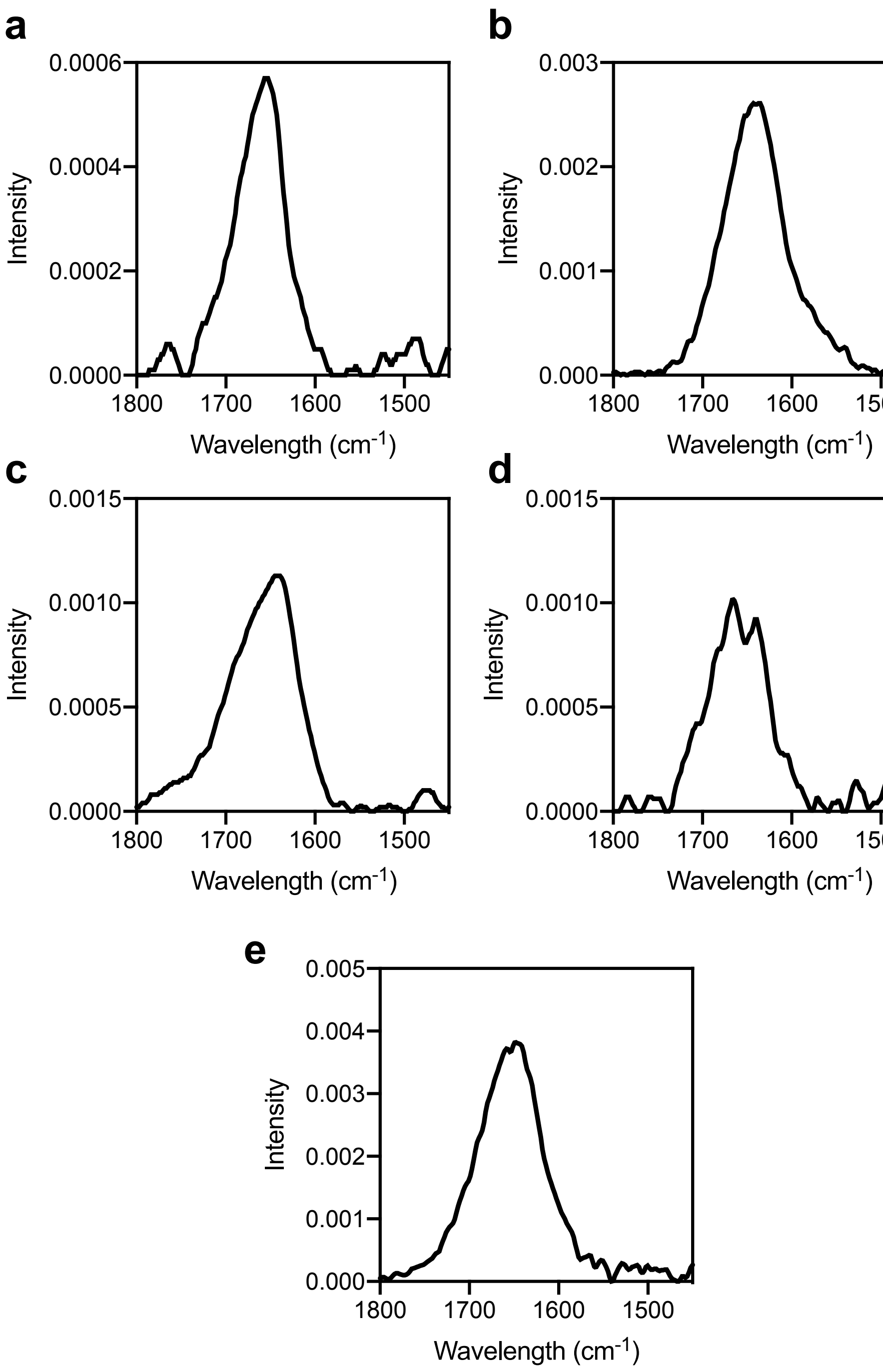


**Figure S7**. FTIR of A$\beta_{1-40}$ fibrils formed in the presence of (a) control, (b) SBA-15, (c) SBA-PFDTS, (d) SBA-PEG, and (e) SiMP.

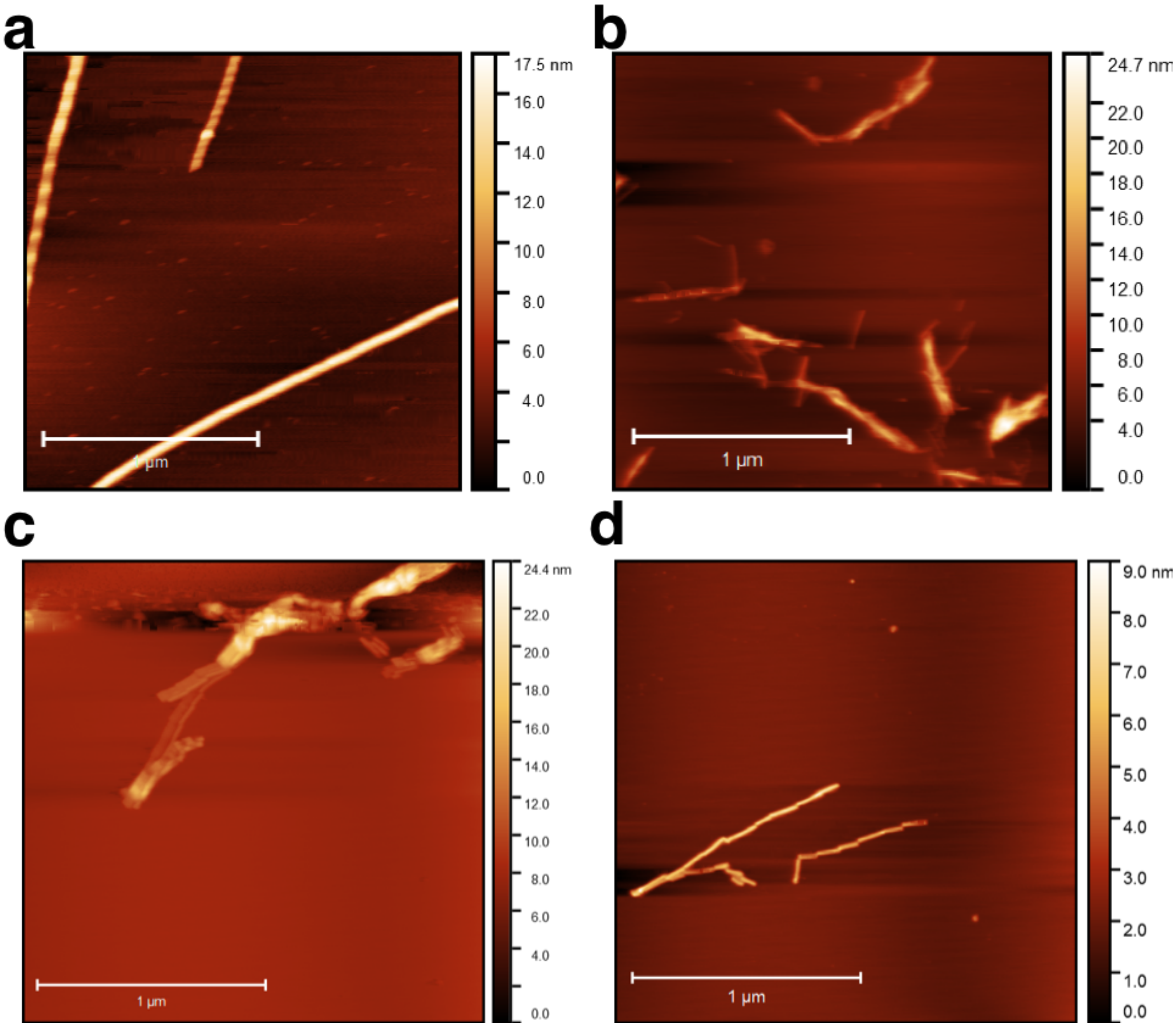


**Figure S8.** AFM of $A\beta_{1\text{-}40}$ fibrils formed in the presence of (a) 10 mM NaPi buffer, (b) SBA, (c) SBA-PEG, and (d) SBA-PFDTS after 24 hr. Fibrils were not detected by AFM for SiMP after 24 hr.

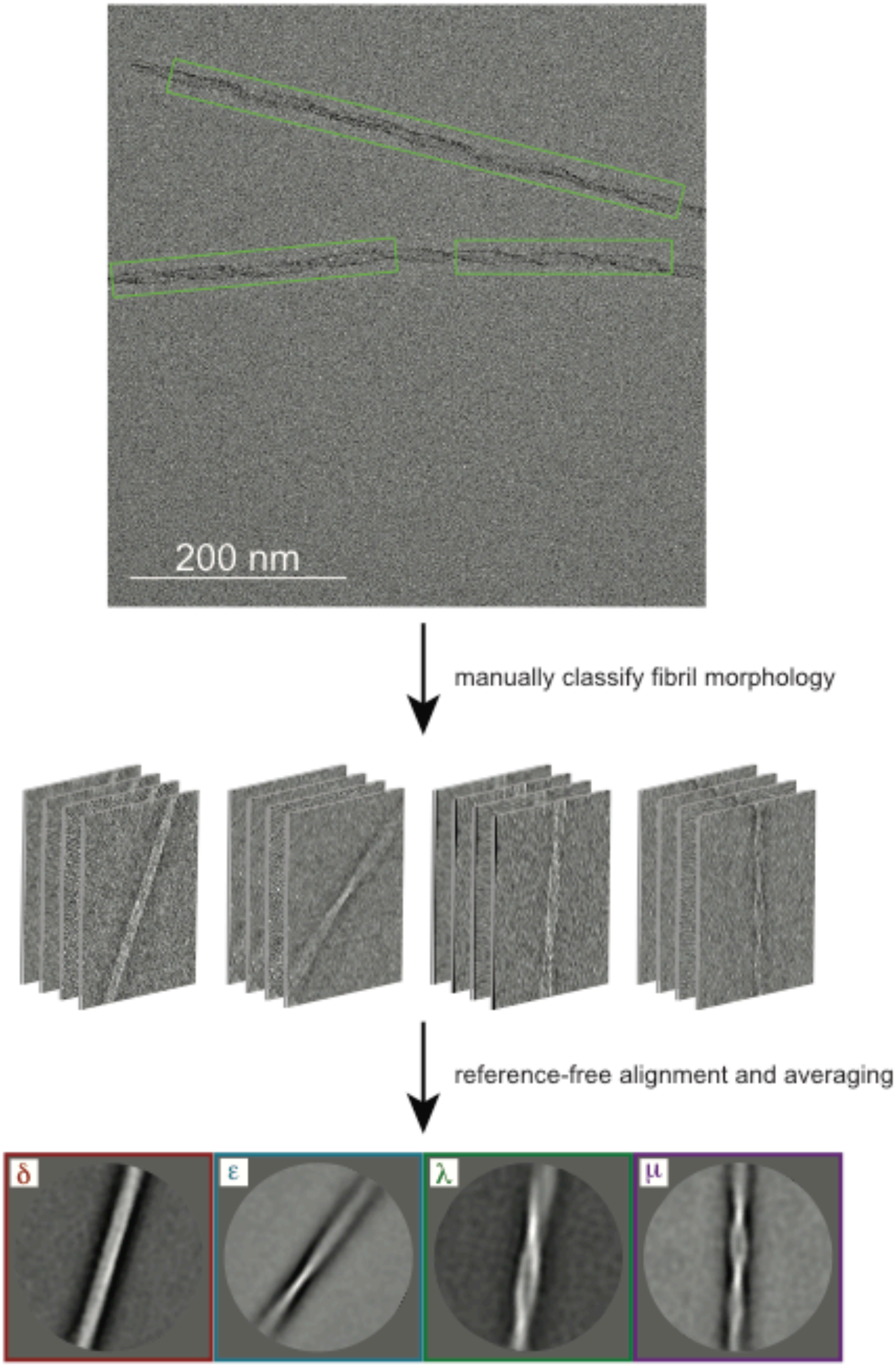


**Figure S9**. Workflow of the production of TEM class-averages of A$\beta_{1\text{-}40}$ fibrils.

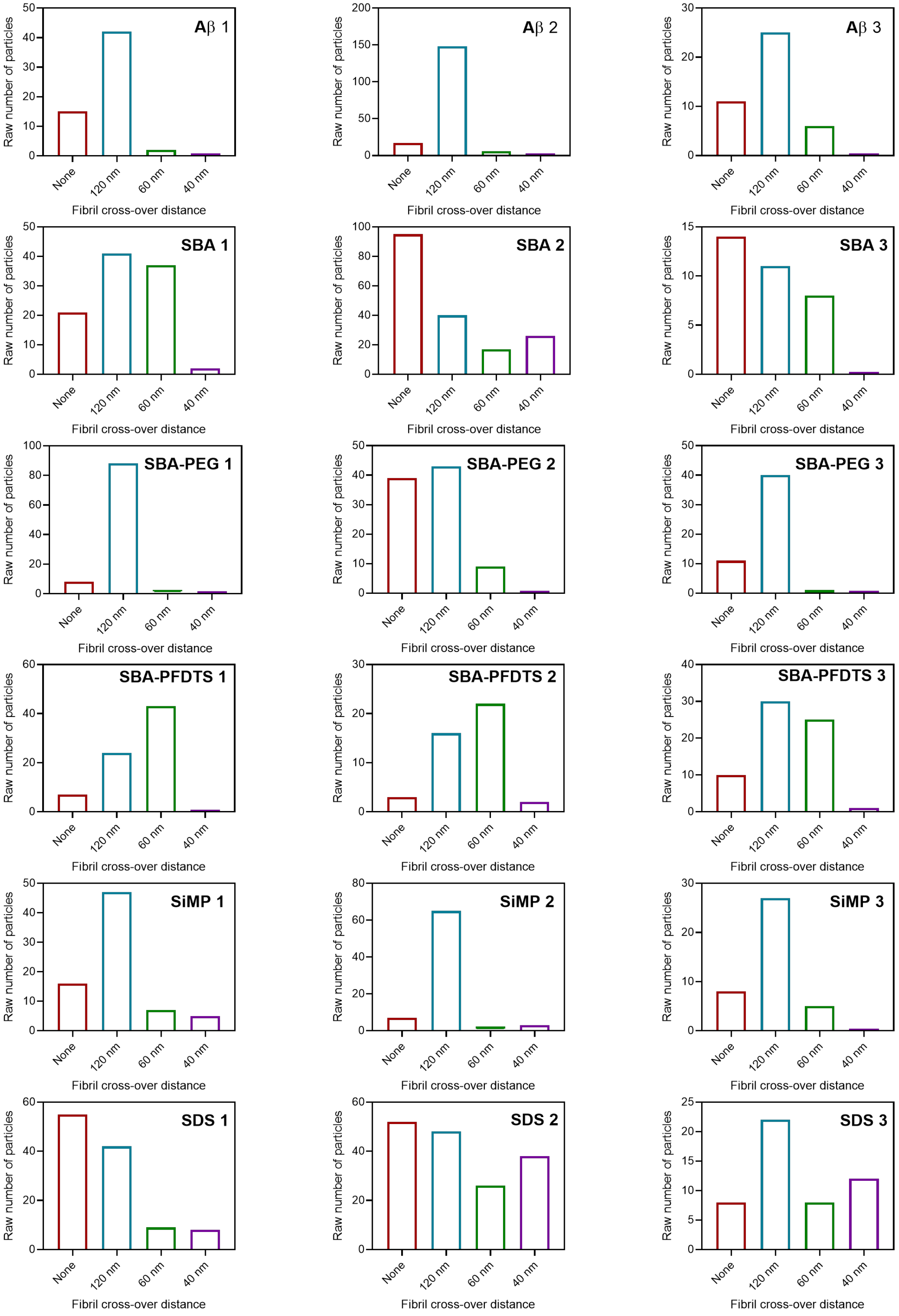


**Figure S10.** Raw counts of $A\beta_{1-40}$ fibril morphologies as measured by negative-stain EM.

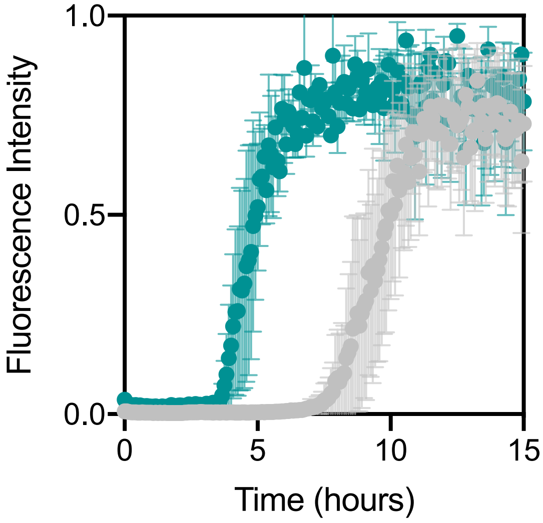


**Figure S11**. Kinetics of fibrilization of A$\beta_{1\text{-}40}$, measured by ThT fluorescence, in the presence of 100 $\mu$M SDS (green) and 10 mM NaPi (grey). Error bars represent one standard deviation. Data show mean ± SD of 3 kinetic runs.

**Table S1**. Surface area and Zeta potential measurements for SBA-15 with functional groups, PFDTS (hydrophobic) and PEG (hydrophilic), attached to the surface.

| | **Surface Area ($m^2/g$)** | **Zeta Potential (mV)** |
|---|---|---|
| SBA-15 | 886 | -20.0 |
| SBA-PEG | 209 | -18.2 |
| SBA-PFDTS | 872 | -20.8 |
| SiMP | 6 | -48.6 |
